# Joint Modeling of Transcriptomic and Morphological Phenotypes for Generative Molecular Design

**DOI:** 10.64898/2026.02.02.703193

**Authors:** Mengbo Wang, Shourya Verma, Vasam Manjveekar Prabantu, Shyaman Jayasundara, Simran Kadadi, Majid Kazemian, Ananth Grama, Nadia Atallah Lanman

## Abstract

**Background:** Effectively translating data on complex cellular responses from transcriptomic and morphological measurements into molecular design remains a significant computational challenge. Existing generative methods operate on single modalities and condition on post-treatment measurements without leveraging paired control-treatment dynamics to capture perturbation effects.

**Results:** We present Pert2Mol, a framework for multi-modal phenotype-to-structure generation that integrates transcriptomic and morphological features from paired control-treatment experiments. Pert2Mol employs bidirectional cross-attention between control and treatment states to capture perturbation dynamics, conditioning a rectified flow transformer that generates molecular structures along straight-line trajectories. We introduce Student-Teacher Self-Representation (SERE) learning to stabilize training in high-dimensional multi-modal spaces. On the Ginkgo Data Platform (GDP) dataset, Pert2Mol achieves Fréchet ChemNet Distance of 4.996 compared to 7.343 for diffusion baselines and 59.114 for transcriptomics-only methods, while maintaining perfect molecular validity and appropriate physicochemical property distributions. The model demonstrates 84.7% scaffold diversity and 12.4 times faster generation than diffusion approaches with deterministic sampling suitable for hypothesis-driven validation.

**Conclusions:** Pert2Mol establishes a new paradigm for linking high-content phenotypic screening data with computational hypothesis generation in drug discovery: joint multi-modal perturbation modeling enables more accurate and structurally diverse molecular design than single-modality or diffusion-based approaches. Code and pretrained models are available at https://github.com/wangmengbo/Pert2Mol.

## 1 Background

Phenotypic drug discovery is re-emerging as a powerful alternative to target-centric strategies, yielding more first-in-class medicines Swinney (2013); Vincent et al. (2022). Modern assays capture transcriptomic and morphological responses to perturbations Chandrasekaran et al. (2021); Bray et al. (2017); however, translating these rich readouts into molecular design remains a challenging inverse problem. Current workflows rely on manual interpretation, creating a bottleneck between phenotypic data and structure–activity connections, particularly since the mapping is inherently many-to-many: different compounds often induce convergent phenotypes Sun et al. (2012), and a single compound may induce multiple phenotypic changes. Rather than a limitation, this redundancy enables scaffold hopping and discovery of novel mechanisms; most approved drugs, in fact, historically emerged from phenotype-first approaches Eder et al. (2014).

Existing AI-based generative methods do not adequately capture the complexity of the molecular design problem. Transcriptome-only models such as MolGene-E use differential expression profiles without paired controls Ohlan et al. (2025), while morphology-based approaches rely solely on imaging Zapata et al. (2023); Tang et al. (2025). No current framework integrates both modalities, despite their complementarity: expression reveals pathway-level changes, while microscopy captures structural phenotypes. Graph-based and text-conditioned models Chang and Ye (2024); Wang et al. (2023) ensure structural validity or semantic alignment but fail on high-dimensional multi-modal perturbation data. Multi-modal conditioning poses additional computational challenges, since the space of perturbations spans thousands of genes and complex morphological features, requiring encoding strategies that preserve interpretability while enabling cross-modal interactions. Diffusion models add inefficiency with large numbers of denoising steps and stochastic outputs misaligned with reproducibility requirements for hypothesis-driven validation Song et al. (2020).

We present Pert2Mol, a framework for multi-modal phenotype-to-structure generation that, to our knowledge, is among the first to jointly integrate transcriptomic and morphological features from paired control–treatment experiments for generative molecular design. By combining these modalities with rectified flow dynamics, Pert2Mol enables efficient, structurally diverse molecule generation guided by complex cellular phenotypes, establishing a paradigm for linking high-content phenotypic screening with computational hypothesis generation in drug discovery. Pert2Mol operates in the latent space of molecular autoencoders, conditioned on integrated transcriptomic and morphological embeddings. A paired transcriptome encoder with cross-attention learns gene-to-gene mappings between control and treatment states, capturing perturbation dynamics beyond differential expression. Morphological features are extracted via a pre-trained image encoder, providing complementary information. These embeddings condition a rectified flow transformer that learns velocity fields along straight-line trajectories from noise to molecular structures, eliminating stochastic sampling and high cost of diffusion models, while enabling deterministic hypothesis generation suitable for validation.

Our contributions include: (i) a rectified flow framework for inverse drug design conditioned on paired multi-modal perturbation signatures; (ii) a transformer architecture that directly models control–treatment perturbation dynamics by fusing transcriptomic and morphological signals; and (iii) Student-Teacher Self-Representation Learning (SERE), where an Exponential Moving Average (EMA) teacher supervises student representations across depths, stabilizing training in high-dimensional multi-modal spaces.

### 1.1 Generative Models for Molecular Design

Generative models for molecular design have advanced from Variational Autoencoders (VAEs) and Recurrent Neural Networks (RNNs) Gómez-Bombarelli et al. (2018) to graph- and flow-based models such as GraphNVP and MoFlow Madhawa et al. (2019); Zang and Wang (2020), which ensure structural validity and latent invertibility. More recently, transformers and diffusion models have improved controllability; e.g., Graph Diffusion Transformers Liu et al. (2024a) integrate property encoders with a transformer denoiser for multi-conditional generation across polymers and small molecules. While these approaches optimize chemical and physicochemical properties (e.g., solubility, synthetic accessibility), they often do not incorporate biological conditioning (transcriptomic, morphological) or perturbation dynamics. Multi-property inverse design has been explored using hierarchical VAEs, Gaussian mixture latent spaces, and encoder-decoder architectures Jin et al. (2018), addressing constraints such as synthetic score and gas permeability, but relying on static property vectors rather than high-dimensional biological readouts or paired control-treatment states.

### 1.2 Perturbation-Conditioned Molecular Design

Perturbation conditioned molecular design incorporates biological data more directly. MolGene-E Ohlan et al. (2025) harmonizes bulk and single-cell transcriptomics with contrastive learning to generate molecules from perturbation-induced profiles. GexMolGen Cheng et al. (2024) produces hit-like molecules guided by gene expression signatures, and GxVAEs Li and Yamanishi (2024) predict responses from transcriptomic profiles. However, these methods are largely unimodal and focus on processed differential or post-treatment signatures rather than paired control-treatment modeling or joint transcriptome-morphology conditioning.

### 1.3 Flow Matching and Biological Conditioning

Flow matching constructs deterministic generative paths and efficient sampling compared to diffusion. GraphNVP Madhawa et al. (2019) and MoFlow Zang and Wang (2020) yield invertible latent representations, but flow-based frameworks conditioned on multi-modal biological data (gene-morphology) under paired perturbation control remain unexplored. Our work addresses this gap by integrating latent rectified flow with multi-modal condition encoders to enable efficient, deterministic molecule generation from biological perturbation data.

## 2 Results

We compared Pert2Mol against two baselines and ablation variants on the GDP Model and Biologics (2025) test set. TransGEM Liu et al. (2024b) was trained on GDP for [[*E*]] epochs using the authors’ official implementation, hyperparameters, and tenfold binary gene-expression encoding, on an NVIDIA A30 GPU. Its training budget is smaller than Pert2Mol’s, so its results are reported as a reference point for transcriptomics-only conditioning rather than as a compute-matched comparison. We randomly selected 1,000 test samples and generated molecules using beam-search decoding with *k* = 10 beams, seq num = 100 sequences per sample, and max len = 60 tokens, yielding ∼ 100,000 total molecules. The diffusion baseline employs DDPM architecture with identical encoder components and latent dimensionality as Pert2Mol, trained with 1000 timesteps using cosine noise schedule. MolGene-E and GexMolGen are not included as baselines because both require differential or post-treatment transcriptomic signatures in their native input format and provide no mechanism for paired control–treatment or morphological conditioning, so they cannot be evaluated on the GDP inputs used here without redefining the task.

### 2.1 Molecular Generation

Quantitative comparisons are shown in Figure 3E. TransGEM exhibits severe mode collapse, generating molecules with mean molecular weight 127.6 Da versus 414.7 Da for reference compounds (3.2-fold reduction). Generated structures consist primarily of linear aldehydes, ketones, and simple heterocycles. Among ∼ 100,000 generated molecules, only 117 unique Bemis-Murcko scaffolds appear (0.117% of generated molecules), versus 912 unique scaffolds in the reference set. KL divergences are 19.08 for molecular weight, 7.71 for LogP, and 20.75 for Topological Polar Surface Area (TPSA), with Frechet ChemNet Distance (FCD) 59.1 reflecting distribution failure. This behavior is consistent with a distributional mismatch between binary-encoded expression features and the continuous profiles in GDP; the reduced training budget noted above may also contribute, and we therefore treat this comparison as indicative of the transcriptomics-only setting rather than as an upper bound on the method. The diffusion model achieves FCD 7.343 versus Pert2Mol’s 4.996, despite requiring 12.4 *×* longer generation time per batch: rectified flow’s straight-line trajectories are both more accurate and more efficient here. Removing SERE degrades FCD from 4.996 to 6.809 (36% increase), consistent with its intended role in stabilizing multi-modal training. The RNA-only variant achieves FCD 5.501 versus 5.762 for image-only, so transcriptomic features carry a stronger perturbation signal on their own. The full multi-modal model achieves the best FCD, Quantitative Estimate of Drug-likeness (QED), Lipinski compliance, and drug-likeness, and closely tracks the reference LogP and TPSA distributions; the SERE-disabled variant instead attains marginally higher Tanimoto similarity and lower molecular-weight KL divergence, indicating that SERE trades a small amount of single-property matching for substantially better overall distributional fidelity (FCD) rather than uniformly dominating every individual metric. Pert2Mol maintains perfect validity while achieving QED 0.587 and Lipinski compliance 78.5%. Figure 3C extends these generation metrics to CPG-jump and LINCS datasets.

### 2.2 Drug Repurposing and Retrieval

All repurposing and retrieval results reported are computed on the 39 Ginkgo Data Platform (GDP) Model and Biologics (2025) compounds, which are shared between the training and test partitions (Section 5.5); the test partition therefore evaluates generalization across unseen experimental conditions for previously seen compounds, not generalization to structurally novel drugs.

Pert2Mol closely matches training statistics (Table 1), with property KL divergences 39–99 *×* lower than TransGEM. Generated molecules have mean molecular weight 408.3 Da (reference: 414.7 Da), LogP 2.79 (reference: 2.83), and TPSA 102.1 (reference: 104.7), with standard deviations closely matching reference values. Pert2Mol generates 847 unique Bemis-Murcko scaffolds among 1,000 molecules (84.7% scaffold diversity); note that the reference set contains 912 unique scaffolds across 38,528 molecules, so a direct percentage comparison is not appropriate. The key finding is that generated scaffolds span a broad and novel chemical space rather than collapsing to a few templates. The diffusion baseline produces 712 unique scaffolds, while TransGEM generates only 117 across 100,000 samples.

**Table 1.** Molecular property statistics for generated molecules vs. reference training set (38,528 compounds). Pert2Mol closely matches reference distributions while TransGEM generates simple fragments.

| Method | Molecular Weight (Da) |  | LogP |  | TPSA |  |
| --- | --- | --- | --- | --- | --- | --- |
|  | Mean | Std | Mean | Std | Mean | Std |
| Reference | 414.7 | 115.1 | 2.83 | 1.67 | 104.7 | 41.0 |
| TransGEM | 120.0 | 25.3 | 1.00 | 0.73 | 37.9 | 11.2 |
| **Pert2Mol** | **408.3** | **112.7** | **2.79** | **1.61** | **102.1** | **39.8** |
| *KL Divergence (lower is better):* |  |  |  |  |  |  |
| TransGEM | 19.08 |  | 7.71 |  | 20.75 |  |
| **Pert2Mol** | **0.26** |  | **0.20** |  | **0.21** |  |

Figure 3B demonstrates effective therapeutic candidate identification, with Tanimoto similarity 0.571 on GDP versus 0.498 for diffusion baseline. The RNA-only variant achieves the highest similarity (0.597); transcriptomic signatures appear to carry the primary signal for compound identification, while the full multi-modal framework remains essential for structural generation fidelity. Image-only and SERE-disabled variants achieve reduced performance (0.407 and 0.401): morphological features alone are insufficient, and SERE plays a clear role in producing robust representations. Figure 2 shows representative repurposed molecules for GDP test perturbations across different conditioning variants. Compounds span diverse mechanisms including mTOR inhibitor Torin2, topoisomerase inhibitor S-Camptothecin, tubulin inhibitors Nocodazole and Colchicine, and proteasome inhibitor Bortezomib.

**Fig. 1.**
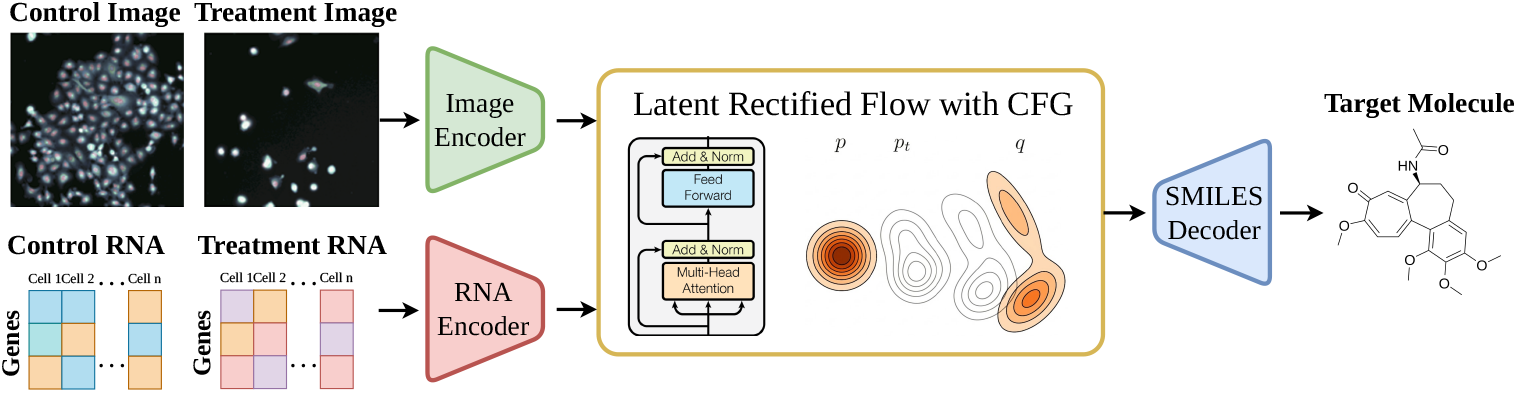
Perturbation guided drug molecule design through Pert2Mol.

**Fig. 2.**
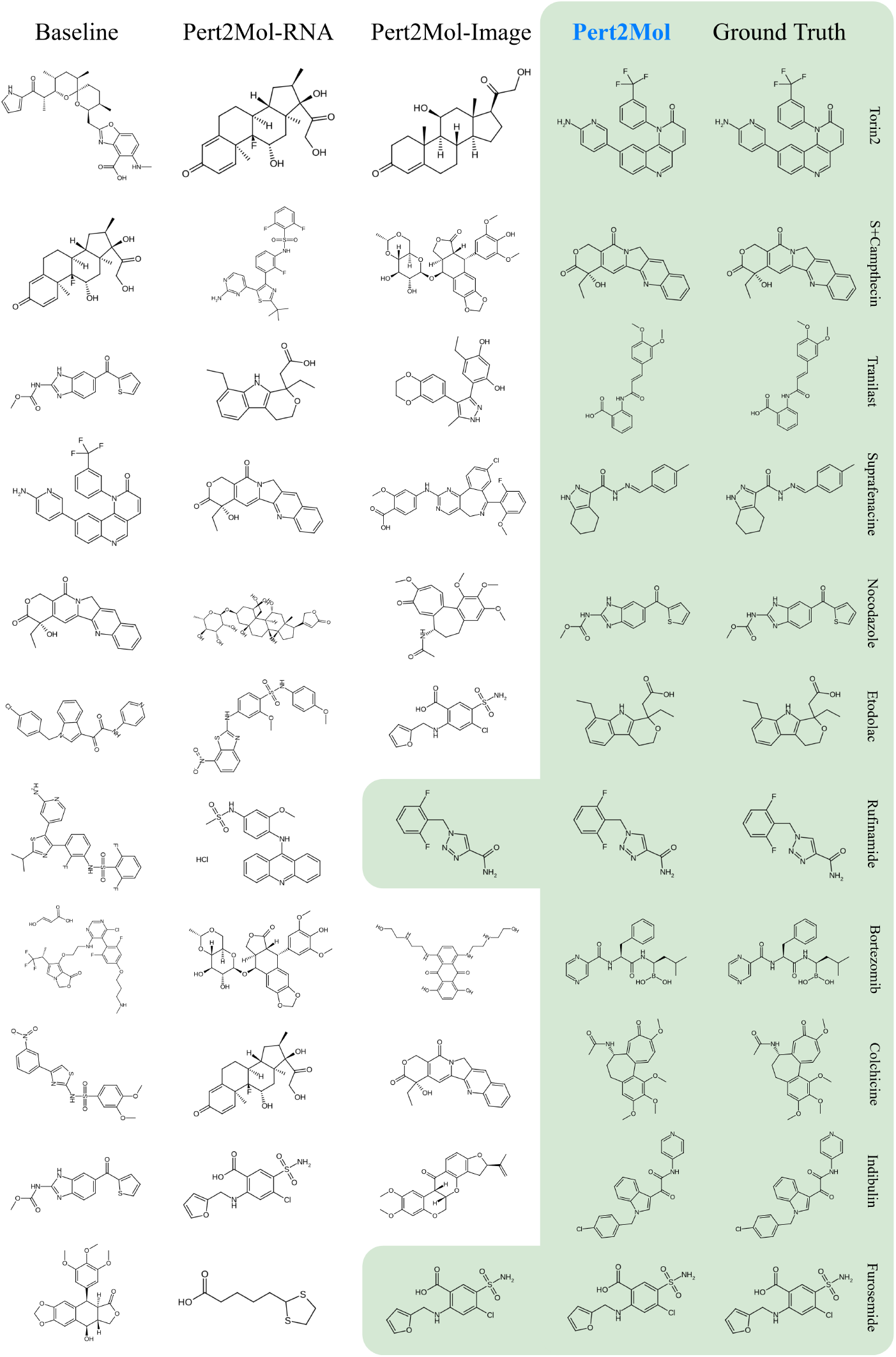
Molecule repurposing results for given control-treatment RNA-image input for different drug examples from GDP dataset. Pert2Mol is compared against diffusion baseline along with RNA-only and image-only models.

**Fig. 3.**
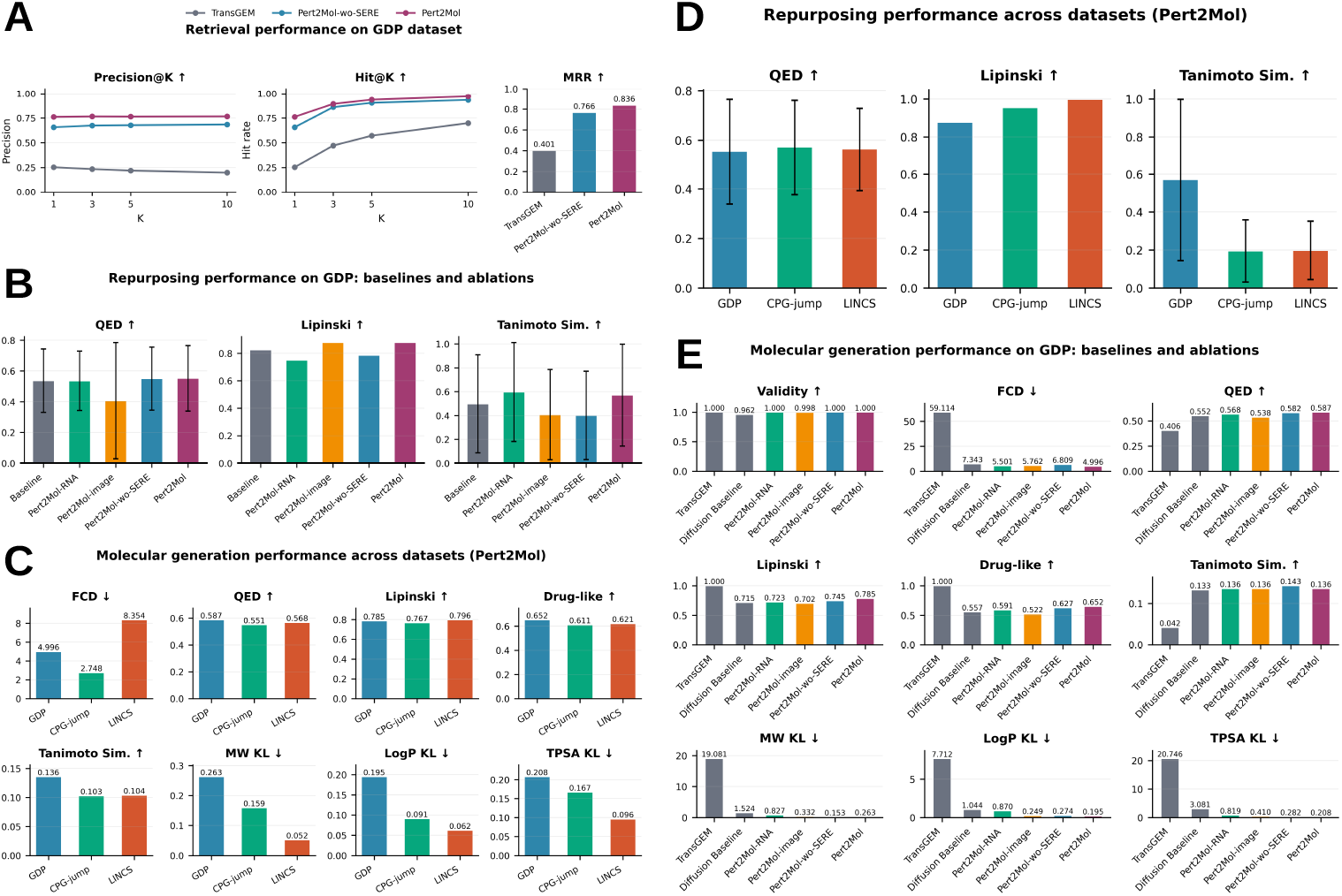
Quantitative evaluation of Pert2Mol on compound retrieval, drug repurposing, and molecular generation. (A) Compound retrieval performance on the GDP training set for TransGEM, Pert2Mol-wo-SERE, and Pert2Mol, reported as Precision@K and Hit@K (K = 1, 3, 5, 10) and Mean Reciprocal Rank (MRR). (B) Drug repurposing performance on the GDP dataset for the baseline and Pert2Mol ablation variants (RNA-only, image-only, without SERE), evaluated by QED, Lipinski compliance, and Tanimoto similarity to ground-truth compounds. (C) Molecular generation performance of the full Pert2Mol model across the GDP, CPG-jump, and LINCS datasets, evaluated by FCD, QED, Lipinski compliance, drug-likeness, Tanimoto similarity, and KL divergence of molecular weight, LogP, and TPSA. (D) Drug repurposing performance of Pert2Mol across the GDP, CPG-jump, and LINCS datasets for QED, Lipinski compliance, and Tanimoto similarity. (E) Molecular generation performance on the GDP dataset for TransGEM, the diffusion baseline, and Pert2Mol ablation variants across the same nine metrics as in (C). Error bars in (B) and (D) denote standard deviation. Arrows next to metric names indicate the direction of improved performance.

We evaluate performance on external unimodal datasets shown in Figure 3D. While GDP achieves 0.571 similarity, the image-only CPG-jump and transcriptomics-only LINCS datasets achieve only 0.195 and 0.198; missing modalities and experimental domain shifts compound to reduce accuracy here. Despite lower retrieval accuracy, the model maintains high validity and Lipinski compliance (95.5% and 99.8%) on external datasets. Figure 3A shows compound retrieval on the GDP training set (evaluated on augmented training samples to test whether the encoder maps each perturbation to its correct location in the drug latent space, rather than to hold out generalization), where Pert2Mol achieves Precision@1 0.7610 and Hit@1 0.7610, versus 0.6560 for SERE-disabled variant. Performance scales to Precision@10 0.7659 and Hit@10 0.9690. Mean Reciprocal Rank increases from 0.7656 without SERE to 0.8364, indicating that representation alignment sharpens the perturbation-to-compound mapping.

### 2.3 Target Engagement Validation: Docking and Transcriptional Pathway Overlap

#### Structure-based docking

To further evaluate whether generated molecules preserve target-specific binding properties beyond global structural similarity metrics, we performed structure-based docking against protein targets associated with the ground-truth compounds. For each reference drug, a primary biological target was identified from literature and pharmacological annotations. Protein structures were obtained using AlphaFold, and molecular docking was performed using AutoDock Vina Trott and Olson (2010). Each ground-truth compound was docked against its corresponding target receptor, and generated molecules conditioned on the perturbation profile of that compound were evaluated against the same cognate receptor.

Ground-truth compounds exhibited a broad range of docking affinities across targets, with several (e.g., Torin2, Corticosterone, Mitoxantrone) showing strongly favorable binding energies. Across the seven compounds retained for quantitative analysis, [[*n*]] generated molecules were docked per cognate receptor, and [[*p*]]% achieved binding affinities at least as favorable as the corresponding ground-truth compound, with a median affinity difference of [[*d*]] kcal/mol (Figure 4D). In particular, for compounds with weak ground-truth binding such as Rotenone (GT affinity ≈ − 1.5 kcal/mol), the majority of generated molecules achieved substantially more favorable docking scores: perturbation-conditioned generation can recover target-compatible chemical scaffolds even when the reference compound itself is a weak binder. Five compounds (BRD3308, Podophyllotoxin, Nocodazole, Colchicine, Suprafenacine) had positive ground-truth docking scores against their annotated targets and were excluded from the quantitative heatmap analysis. The 5-of-12 exclusion rate most likely reflects known limitations of rigid, single-chain AlphaFold-model docking with AutoDock Vina Scardino et al. (2023) Karelina et al. (2023) Pantsar and Poso (2018); e.g., missing cofactors, multimeric assemblies, or allosteric/covalent binding modes not captured by this pipeline, rather than these compounds genuinely failing to engage their reported targets. We therefore treat the remaining seven-compound panel as the more reliable basis for quantitative comparison and interpret the docking results as a supporting, rather than definitive, line of evidence.

**Fig. 4.**
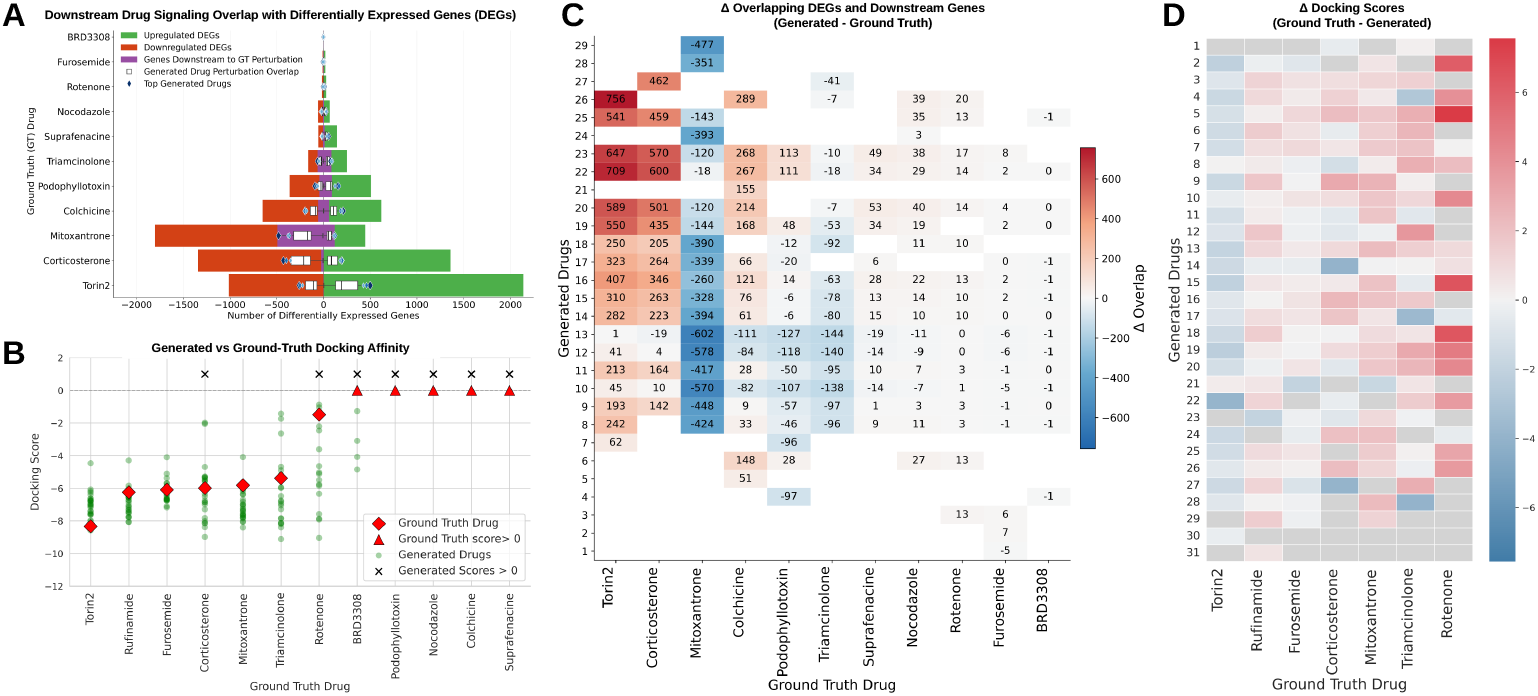
Differential gene expression, target-downstream pathway overlap, and target-specific docking comparison between ground-truth compounds and generated molecules. (A) Differentially expressed genes (DEGs) and target-downstream pathway overlap for ground-truth perturbations. For each ground-truth compound (rows, sorted by total DEG count), green and orange bars show the number of up- and down-regulated DEGs, respectively. The purple line shows the overlap between the compound’s DrugCLIP-predicted target-downstream gene set and its own DEGs (ground-truth overlap). White box plots show the distribution of the corresponding overlap for the target-downstream gene set of molecules generated under that compound’s perturbation profile, evaluated against the same DEG set; blue diamonds mark the top generated candidates. (B) Docking affinity scores for all cognate ligand–receptor pairs, ordered by ground-truth affinity. Green circles denote docking affinities of generated molecules conditioned on the perturbation profile of each compound; red diamonds denote the ground-truth compound. Ground-truth compounds and generated molecules with positive docking scores (unfavorable predicted binding) are shown as red triangles and black crosses, respectively. (C) Heatmap of Δ overlap (generated molecule’s target-downstream overlap minus the ground-truth compound’s own overlap) for each generated molecule (rows, numbered) evaluated against every ground-truth compound’s DEG set (columns), not only the one it was conditioned on. Red indicates the generated molecule’s target-downstream genes overlap the column compound’s DEGs more strongly than the ground-truth compound’s own target does; blue indicates weaker overlap. Blank cells were not evaluated. (D) Heatmap of the difference between ground-truth and generated docking affinity (ground-truth affinity minus generated affinity) for the seven compounds whose ground-truth docking scores were favorable (*<* 0). Positive values (red) indicate generated molecules with stronger predicted binding than the ground-truth compound; negative values (blue) indicate weaker predicted binding. Grey cells correspond to molecules not evaluated against a given receptor.

Figure 4 summarizes the docking analysis across all matched target–ligand systems. Panel B shows the distribution of docking affinities across cognate receptors, sorted by ground-truth affinity; panel D highlights per-molecule affinity differences for the seven compounds with favorable ground-truth docking.

#### Downstream transcriptional pathway overlap

As a complementary, targetagnostic measure of mechanistic plausibility, we asked whether the genes downstream of a compound’s predicted target, in a curated signaling network, overlap with the differentially expressed genes (DEGs) actually measured after treatment. Protein targets for both ground-truth and generated compounds were identified using genome-wide contrastive virtual screening Gao et al. (2023), and downstream genes were obtained by one-hop traversal of the OmniPath signaling network Türei et al. (2016) from each predicted target. For each ground-truth compound, this downstream gene set was intersected with the DEGs measured in the corresponding perturbation experiment. For each generated molecule, the same procedure was applied to its own predicted target and downstream gene set, and the resulting overlap was compared against the DEG sets of every ground-truth compound in the panel, not only the one it was conditioned on, yielding a full generated-molecule *×* ground-truth-compound overlap matrix.

Figure 4A shows that both the number of DEGs (1 for BRD3308 to 3,150 for Torin2) and the size of the predicted target’s downstream gene set (46 genes for Torin2 to 4,385 for Triamcinolone) vary by more than two orders of magnitude across ground-truth compounds. Because raw overlap counts scale with both quantities, they are not directly comparable across compounds; we therefore normalize each ground-truth overlap by its downstream-set size and compare it to the compound’s genome-wide DEG rate (DEGs / ∼ 20,000 protein-coding genes) as a chance baseline. By this measure, Mitoxantrone, Colchicine, Podophyllotoxin, Triamcinolone, Suprafenacine, and Nocodazole show 1.5–2.4-fold enrichment of DEGs among their predicted target’s downstream genes relative to chance, so the predicted target’s network neighbor-hood does appear to capture a meaningful, if partial, share of the true transcriptional response. Torin2 is a clear outlier: only 4 of its 46 downstream genes (8.7%) are DEGs, below the 15.8% expected by chance given its 3,150 DEGs, and its predicted target also had the lowest screening confidence in the panel, suggesting that DrugCLIP’s target assignment for Torin2 is unreliable rather than that Torin2 lacks a coherent downstream pathway; Torin2’s ground-truth overlap value is therefore an artificially weak baseline for comparison.

The distinction between compounds with above- and below-chance ground-truth overlap matters for interpreting Figure 4C: several molecules generated under other perturbation profiles (e.g., Corticosterone-conditioned candidates) show large positive Δ overlap specifically in the Torin2 column, exceeding Corticosterone’s own column by a similar margin. Given that Torin2’s ground-truth baseline is deflated by a low-confidence, narrow (46-gene) target assignment, almost any generated molecule whose predicted target has an average-sized downstream footprint will numerically outscore it; this pattern is more parsimoniously explained by a weak reference baseline than by genuine cross-drug mechanistic similarity. By contrast, for compounds whose ground-truth overlap already exceeds chance (Colchicine, Podophyllotoxin, Mitoxantrone, Triamcinolone), Δ overlap in the corresponding column is a more trustworthy indicator, and here the pattern is better grounded: overlap for molecules generated under Colchicine, Podophyllotoxin, and Furosemide conditioning is concentrated in the correct column and small in magnitude rather than spread broadly across the panel, and overlap against Mitoxantrone’s DEG set remains negative for nearly all molecules not conditioned on it. Target-downstream overlap therefore provides genuine, if modality-dependent, mechanistic support for several perturbation-conditioned generations; comparisons involving compounds with low-confidence or unusually narrow target predictions, such as Torin2 in this panel, should be discounted rather than read as evidence of superior recovery.

## 3 Discussion

Pert2Mol is an effective framework for multi-modal phenotype-to-structure generation that integrates transcriptomic and morphological features from paired control-treatment experiments. Our latent rectified flow method consistently outperforms diffusion-based approaches across drug discovery metrics while maintaining perfect molecular validity. The full multi-modal model achieves an FCD of 4.996 compared to 7.343 for the diffusion baseline and 59.114 for TransGEM, a substantial improvement in capturing the training distribution’s chemical space.

Rectified flow offers deterministic sampling with 12.4 times faster generation than diffusion models while maintaining superior sample quality. The straight-line velocity field enables efficient sampling in fewer steps without complex noise scheduling, addressing reproducibility requirements for hypothesis-driven validation. Property distribution analysis confirms faithful preservation of physicochemical characteristics, with KL divergences of 0.263 for molecular weight, 0.195 for LogP, and 0.208 for TPSA. Generated molecules exhibit 84.7% scaffold diversity (847 of 1,000 unique Bemis-Murcko scaffolds), compared to 71.2% for the diffusion baseline under the same protocol, consistent with genuine exploration of chemical space rather than mode collapse; as noted above, a direct percentage comparison to the reference training set’s scaffold diversity is not meaningful given its far larger size (912 unique scaffolds across 38,528 compounds).

SERE provides training stability crucial for high-dimensional multi-modal spaces, with ablation studies showing FCD degradation from 4.996 to 6.809 when SERE is removed. The bidirectional cross-attention mechanism enables direct learning from paired experimental data. Notably, our ablation analysis reveals a task-dependent modality preference: while the RNA-only variant excels at compound identification, the full multi-modal integration is required to maximize generative fidelity (FCD 4.996 vs. 5.501 for RNA-only): transcriptomics appears to provide a distinct “fingerprint” for retrieval, while the complementary morphological information is essential for constraining the generative flow to the correct structural manifold.

Cross-dataset evaluation highlights the challenge of generalization to unimodal contexts. Performance varies substantially, with FCD scores of 4.996 on GDP (which is biased toward kinase and HDAC inhibitors with molecular weights between 200–600 Da) compared to 2.748 on the image-only CPG-jump and 8.354 on the transcriptomics-only LINCS dataset. Because GDP, CPG-jump, and LINCS differ in chemical composition, assay technology, and available modalities all at once, these three numbers are not a controlled test of whether multi-modal conditioning outperforms unimodal conditioning; that comparison is instead made within GDP, where the paired RNA-only and image-only ablations (Figure 3E) show the full multi-modal model outperforming either single modality on the same compounds and the same encoder-decoder pipeline. The lower FCD on CPG-jump reflects that dataset’s narrower, less complex chemical space rather than an advantage of image-only conditioning per se, and the higher FCD on LINCS reflects both a missing modality and the added domain shift of a different profiling technology; the TransGEM and diffusion baselines were trained and evaluated only on GDP, so their FCD values are not directly comparable to Pert2Mol’s CPG-jump or LINCS results. Unlike the GDP results (Section 5.5), the CPG-jump and LINCS evaluations involve compounds never seen during training, so they test generalization to structurally novel drugs rather than to unseen conditions of known compounds. Despite these limitations, Pert2Mol demonstrates that multi-modal phenotype-to-structure generation is feasible, offering a systematic approach for translating high-content screening data into molecular hypotheses for drug discovery.

We acknowledge that Tanimoto similarity to ground-truth drugs is an imperfect criterion in phenotypic discovery, where scaffold hopping is often desirable. The higher repurposing similarity of the RNA-only variant (0.597 vs. 0.571) reflects transcriptomics’ stronger direct signal for compound identity; the full multi-modal model trades marginal similarity for better generative fidelity (FCD 4.996 vs. 5.501). Future work should incorporate mechanism-of-action enrichment analyses for stronger biological validation.

## 4 Conclusions

We introduced Pert2Mol, a multi-modal generative framework that translates paired control-treatment transcriptomic and morphological phenotypes directly into drug-like molecular structures. By combining bidirectional cross-attention perturbation encoders, rectified flow generation, and SERE representation learning, Pert2Mol achieves state-of-the-art chemical distribution fidelity, scaffold diversity, and generation efficiency on the GDP benchmark while generalizing to external imaging (CPG-jump) and transcriptomic (LINCS) datasets. These results demonstrate the feasibility of systematic, multi-modal phenotype-to-structure generation and open a path toward data-driven hypothesis generation in phenotypic drug discovery.

## 5 Methods

We formulate drug perturbation prediction as a conditional generative modeling problem, where the model learns to generate molecular structures from multi-modal biological context. Let **x** ∈ ℝ^*D*^ be a molecular structure in latent space, let **c**_img_ ∈ ℝ^4*×H×W*^ be control and treatment microscopy images, and **c**_rna_ ∈ ℝ^*G*^ be gene expression profiles with *G* genes. The objective is to learn conditional distribution *p*(**x** | **c**_img_, **c**_rna_) that captures a relationship between biological perturbation context and molecular structure. Figure 5 presents the high-level architecture of Pert2Mol.

**Fig. 5.**
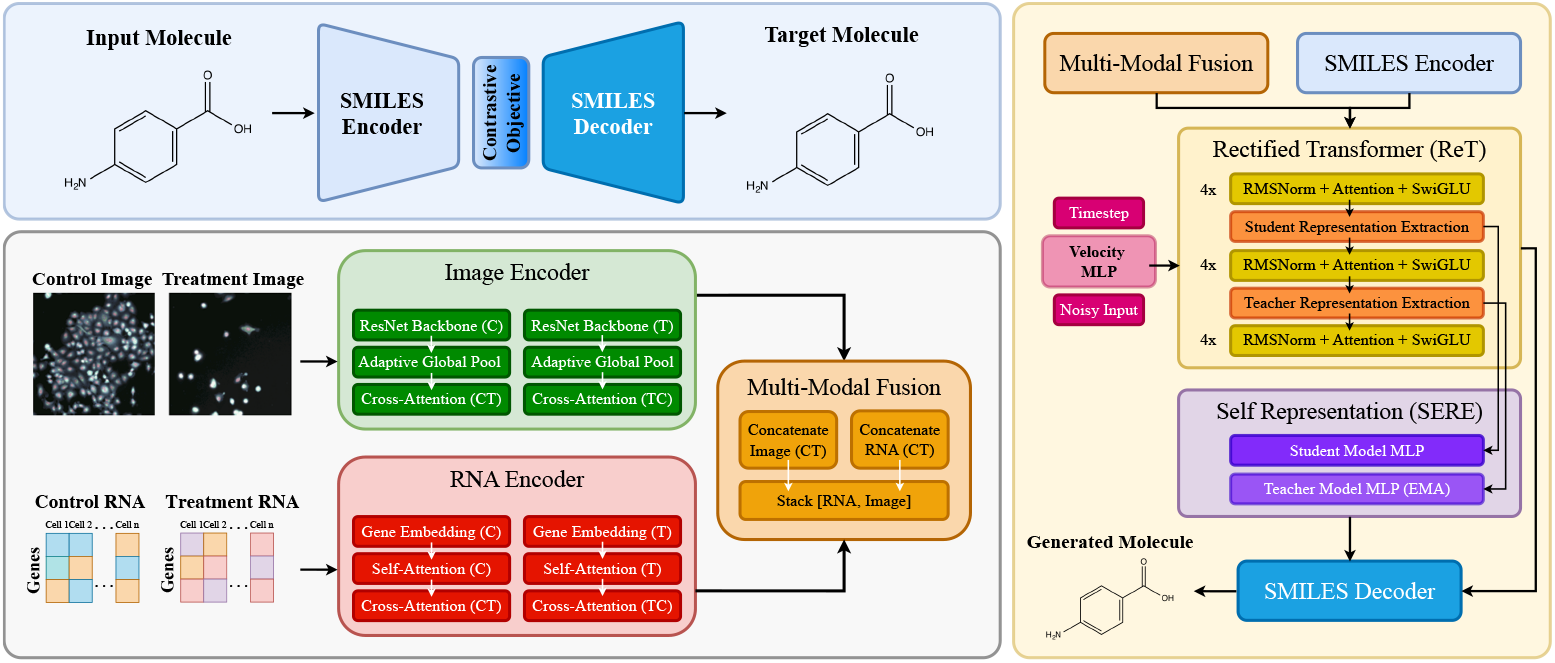
Overview of the Pert2Mol architecture. **(Top left)** A SMILES autoencoder embeds molecules, with a contrastive objective aligning the encoder latent space to the perturbation representation. **(Bottom left)** Paired control and treatment inputs are encoded separately: cell images through a ResNet backbone with adaptive global pooling and bidirectional cross-attention (C → T, T → C), and bulk gene expression through gene embeddings, self-attention, and the same bidirectional cross-attention. The two modalities are concatenated and stacked into a joint multi-modal fusion token sequence. **(Right)** A Rectified Transformer (ReT) conditioned on the fused perturbation representation, the diffusion timestep, and a velocity MLP denoises the noisy SMILES embedding through blocks of RMSNorm, attention, and SwiGLU. Student and teacher representations extracted at intermediate depths feed the Self Representation (SERE) module, where a teacher MLP updated by EMA supervises the student MLP. The resulting embedding is decoded into the generated molecule. During training the model receives cell images, bulk gene expression, and SMILES embeddings; at inference only cell images and bulk gene expression are required.

### 5.1 Multi-modal Conditioning

#### Image encoding

We encode paired control and treatment CellPainting images using a frozen pre-trained image backbone *f*_img_ that extracts spatial patch-level representations. For each condition, the backbone produces a sequence of patch tokens:

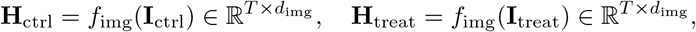

where *T* is the number of patches and *d*_img_ is the backbone embedding dimension. A lightweight trainable bidirectional cross-attention adapter then captures treatment-induced morphological changes:

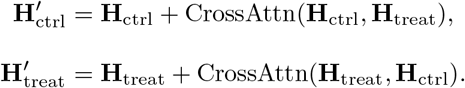

Each adapted sequence is compressed via attention pooling and a linear projection to yield 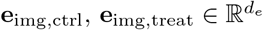:

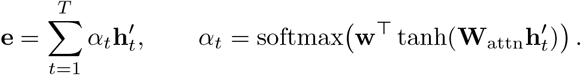

In this work, we instantiate *f*_img_ with the OpenPhenom CA-MAE ViT-S/16 Kraus et al. (2024) (*d*_img_ = 384, *d*_*e*_ = 128), pre-trained by Recursion Pharmaceuticals on large-scale CellPainting screens. Only the adapter and projection layers are trained.

#### RNA expression encoding

Transcriptomic profiles are encoded with a frozen pre-trained RNA backbone *f*_rna_ that maps gene expression vectors to per-gene contextual representations. We subsample to the top *G*^*′*^ highly variable genes to control sequence length:

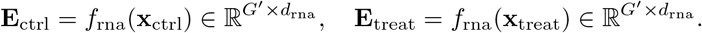

A trainable bidirectional cross-attention adapter then models perturbation dynamics, explicitly capturing gene-to-gene relationships between control and treatment states beyond simple differential profiles:

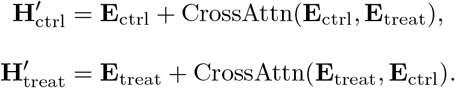

Attention-weighted pooling and separate linear projectors produce condition-level embeddings 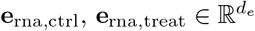:

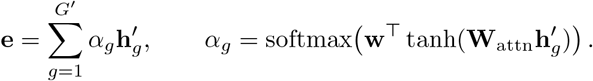

In this work, we instantiate *f*_rna_ with scFoundation (xTrimoGene) Hao et al. (2024) (*d*_rna_ = 768, *G*^*′*^ = 2,000, *d*_*e*_ = 128), a transformer pre-trained on 50 million human single cells that accepts log1p-normalized expression and processes only non-zero gene tokens. Only the adapter and projection layers are trained.

#### Multi-modal fusion

Image and RNA features for each condition are concatenated and stacked to produce the final conditioning tensor:

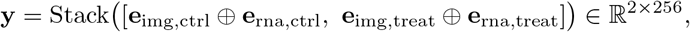

where ⊕ denotes concatenation. This structured representation preserves relationships between conditions while providing rich multi-modal context to the generator.

#### Molecular representation

Molecular structures are represented using a pre-trained BERT-based autoencoder Liu et al. (2019) that maps SMILES strings to continuous latent representations similar to Chang and Ye (2024). Tokenization uses learned molecular motifs (regex tokenization) instead of character-level tokens to capture chemically meaningful substructures. A frozen pre-trained BERT encoder processes tokenized sequences, and a trainable linear compression layer reduces BERT outputs from 768 to 64 dimensions, producing a compact molecular latent representation:

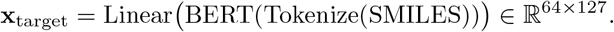

### 5.2 Generative Modeling

We adopt rectified flow Liu et al. (2022) as the generative framework for its training stability and sampling efficiency relative to traditional diffusion models. Rectified flow parameterizes straight-line paths between noise and data distributions. Given a noise sample **x**_0_ ∼ *N* (0, **I**) and data **x**_1_ = **x**_target_, the interpolating path is **x**_*t*_ = (1 − *t*)**x**_0_ + *t***x**_1_, *t* ∈ [0, 1], and its instantaneous velocity is:

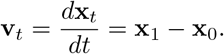

The model is trained to predict this velocity field **v**_*θ*_(**x**_*t*_, *t*, **y**) ≈ **v**_*t*_ with the rectified flow objective:

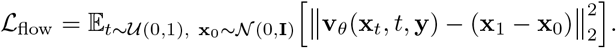

#### Rectified transformer architecture

Our core generative model is a transformer adapted to molecular generation with multi-modal conditioning. Noisy molecular representations are linearly projected and augmented with learnable positional embeddings: 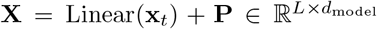, and timesteps are embedded via a sinusoidal encoding passed through an MLP: 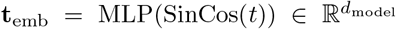. Conditioning information is processed along two pathways:

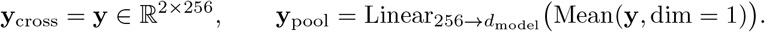

The conditioning signal used for adaptive normalization is **c** = **t**_emb_ + **y**_pool_. Each transformer block implements Adaptive Layer Normalization (AdaLN):

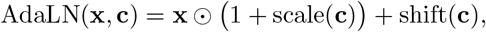

where scale(**c**), shift(**c**) = Linear(SiLU(**c**)). Each block applies self-attention, cross-attention to the conditioning embedding, and a feed-forward update using SwiGLU nonlinearity Shazeer (2020):

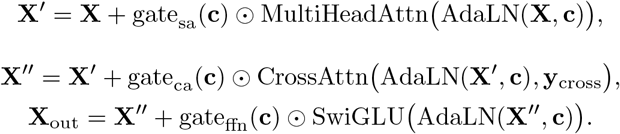

The SwiGLU activation is defined as SwiGLU(**x**) = SiLU(**xW**_1_) ⊙ (**xW**_2_)**W**_3_.

#### Student-Teacher Self-Representation (SERE)

To improve training stability and sample quality, we introduce SERE. The SERE teacher model is maintained as an exponential moving average (EMA) of the student parameters: *θ*_teacher_ ← *β θ*_teacher_ +(1 − *β*) *θ*_student_. For a given training example at timestep *t*, the student network processes the noisy sample **x**_*t*_ as usual, while the teacher network (using the EMA weights) processes the same sample at a less noisy effective timestep *t* − Δ*t* (clamped to ≥ 0); the noise-level difference between student and teacher therefore comes from this timestep shift, not from network depth. Intermediate representations are then extracted from a shallower layer of the student’s forward pass (e.g., **h**_student_ = **X**_layer-4_) and a deeper layer of the teacher’s forward pass (e.g., 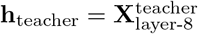): because the teacher operates on a less noisy input, its later-layer representation is a more reliable self-supervisory target than the student’s own forward pass could provide at the same depth. The SERE loss aligns representations using a projection head:

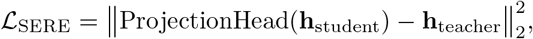

with the projection head defined as:

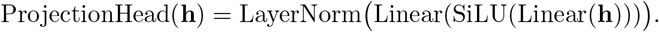

The total training objective combines the rectified flow loss and the SERE loss:

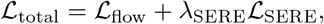

where *λ*_SERE_ = 0.1 balances the contribution of representation alignment.

#### Training setup

Training proceeds in two phases to leverage pre-trained foundation model representations.

*Encoder pretraining*. The cross-attention adapters and projection MLPs of both encoders are jointly pre-trained with all backbone weights frozen. The objective is:

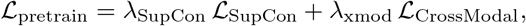

where ℒ_SupCon_ is Supervised Contrastive Loss Khosla et al. (2020) applied to RNA embeddings with drug identity as labels, pulling together same-drug representations across cell lines and pushing apart different drugs. ℒ_CrossModal_ is a soft-label bidirectional InfoNCE loss that aligns pseudo-paired RNA and image embeddings, assigning soft targets of 1.0 for exact condition matches (same cell line, drug, dose, timepoint) and 0.5 for same-drug-different-condition pairs.

*Flow-matching training*. Pre-trained encoder weights are loaded with the foundation model backbones frozen; adapter layers are updated at a 0.1*×* learning rate multiplier relative to the ReT model (*λ*_enc_ = 0.1). For classifier-free guidance Ho and Salimans (2022), modalities are randomly masked during training: 10% RNA-only and 10% image-only inputs, with the remaining 80% fully conditioned; unlike standard classifier-free guidance, the two modalities are never masked simultaneously, so the model never observes a fully unconditional input during training. SMILES strings are augmented via canonical randomization to preserve molecular identity. Optimization uses AdamW (lr=10^−4^, cosine schedule with weight decay), mixed-precision (bfloat16) for memory efficiency, gradient clipping (norm=5.0), and distributed data-parallel training. Two EMA buffers with the same decay rate (*β* = 0.9999) are maintained for distinct roles: one continuously supplies the SERE teacher’s weights throughout training, while the other accumulates independently and is read out only once, after training, for final inference-time parameter averaging.

Sample generation integrates the learned velocity field from noise to data via **x**_*i*+1_ = **x**_*i*_ +Δ*t* · **v**_*θ*_(**x**_*i*_, *t*_*i*_, **y**) using Euler integration with [[*N*]] uniformly spaced steps. Generation throughput was measured on a single NVIDIA H100 GPU at batch size [[*B*]] under identical decoding settings for both models; Pert2Mol requires [[*N*]] network evaluations per sample versus 1000 for the diffusion baseline, giving the 12.4 *×* wall-clock speedup reported in Section 2. Multi-modal classifier-free guidance scales each modality’s contribution independently:

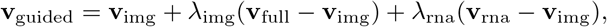

where **v**_full_, **v**_rna_, and **v**_img_ are velocity predictions conditioned on both modalities, RNA-only, and image-only inputs respectively. Guidance is anchored on the image-only prediction **v**_img_ rather than a fully unconditional prediction: because training never masks both modalities simultaneously (above), the model never observes a fully unconditional input, making a null-conditioning anchor out of distribution; the image-only prediction is instead a conditioning regime the model was explicitly trained to handle, and both the RNA-only and full predictions are guided relative to it. Guidance scales were selected by grid search over [[*range*]] on a held-out validation split, minimizing FCD, giving *λ*_rna_ = 5.0 and *λ*_img_ = 4.0, which are used for all reported results. On external datasets only one modality is available, so inference is run in the corresponding single-modality regime the model was explicitly trained to handle under the masking schedule above: the missing modality’s encoder input is dropped and guidance reduces to a single term, using *λ*_img_ = 4.0 for the image-only CPG-jump data and *λ*_rna_ = 5.0 for the transcriptomics-only LINCS data. Latent representations are decoded to SMILES via the pre-trained autoencoder with beam search. The ReT model is a 12-layer transformer (768 hidden dimensions, 16 attention heads, 199M parameters) trained for 300 epochs with batch size 32 on 4 NVIDIA H100 GPUs.

### 5.3 Structure-based Molecular Docking

To evaluate target-specific binding properties of generated molecules, we performed molecular docking using AutoDock Vina Trott and Olson (2010). All docking calculations were conducted using a customized pipeline integrating RDKit Landrum (2016), OpenBabel O’Boyle et al. (2011), and the Vina Python API.

Protein structures corresponding to each ground-truth compound were obtained from the AlphaFold Protein Structure Database Varadi et al. (2022). For each compound, a primary biological target or mechanism-associated protein was selected based on curated literature annotations. Of the 39 GDP compounds, 12 satisfied the inclusion criteria for docking: a single unambiguous primary protein target in the curated annotations, a corresponding human UniProt entry, and an available AlphaFold model of that entry. The remaining compounds were excluded because their annotated mechanism was multi-target, non-protein, or lacked a one-to-one target assignment, and were not evaluated further. Receptor structures were processed by retaining a single protein chain. Hydrogen atoms were added prior to conversion to PDBQT format using OpenBabel.

Ligand structures were generated using RDKit. For each ligand, hydrogen atoms were explicitly added and a 3D conformer was generated using the ETKDG algorithm; molecular geometries were minimized and ligands converted to PDBQT format for docking compatibility. Because AlphaFold models are apo structures without bound ligands, no experimentally defined binding site was imposed. The search box was instead centered on the centroid of the receptor chain’s heavy-atom coordinates, and a cubic search space of 20 *×* 20 *×* 20 Å^3^ was applied identically to every receptor– ligand pair. This site-agnostic protocol places ground-truth and generated ligands under exactly the same search conditions, so affinities are directly comparable within a receptor; absolute affinities are correspondingly interpreted as relative rather than calibrated binding predictions. For each run, binding affinities and ranked generated poses were computed, from which the most favorable (lowest energy) pose was selected as the best docking score for each ligand–receptor pair. All docking calculations were performed using identical parameters across ground-truth and generated molecules to ensure comparability. Ligand–receptor pairs with unfavorable ground-truth docking energies (positive binding affinities) were excluded from downstream quantitative analysis.

### 5.4 Downstream Transcriptional Pathway Overlap Analysis

To complement structure-based docking with a target-agnostic, pathway-level check, we evaluated whether the predicted target of each compound is transcriptionally consistent with its measured perturbation response. Protein targets for both ground-truth and generated compounds were identified by genome-wide contrastive virtual screening against a library of *>*2,000 candidate protein pockets using DrugCLIP Gao et al. (2023); hits with *z*-score ≥ 4.0 were retained as predicted binding partners and mapped to gene identifiers via UniProt. Downstream signaling genes were obtained by one-hop directed traversal, using NetworkX, of the OmniPath signaling network Türei et al. (2016) seeded from each compound’s predicted targets. Differentially expressed genes (DEGs) for each ground-truth compound were called from the corresponding treatment-versus-control experiment (adjusted *p <* 0.05, | log_2_ FC | *>* 1) and split into up- and down-regulated sets. For each ground-truth compound, its down-stream gene set was intersected with its own DEG set to obtain a ground-truth overlap. For each generated molecule, the same downstream gene set was computed from its own DrugCLIP-predicted target and intersected against the DEG sets of every ground-truth compound in the panel (not only the one it was conditioned on), yielding a full generated-molecule *×* ground-truth-compound overlap matrix and, for each cell, a Δ overlap relative to the corresponding ground-truth baseline. Ground-truth compounds lacking matched experimental DEG data (Rufinamide, Amsacrine hydrochloride, Wortmannin) were excluded as reference columns but their conditioned generated molecules were retained as rows, evaluated against the DEG sets of the remaining compounds.

### 5.5 Datasets

We build upon the multi-modal Ginkgo Data Platform (GDP) collection Model and Biologics (2025), which pairs transcriptomic profiles with four-channel fluorescence microscopy of chemically perturbed cell populations. Correspondence across modalities was established through compound identity matching and experimental parameter alignment, with metadata standardized for dose normalization, consistent cell line identifiers, and synchronized timepoints; vehicle (DMSO)-treated samples served as controls. The GDP collection comprises 39 compounds; train/test partitioning was performed at the level of individual experimental conditions (cell line, dose, and time-point combinations), stratified by compound identity, rather than by withholding entire compounds. All 39 compounds therefore appear in both partitions, and retrieval, repurposing, and generation metrics reported on GDP measure generalization across experimental conditions for previously seen compounds, not generalization to structurally novel drugs. Transcriptomic data were TPM-normalized, log1p transformed, and reduced to the top 2000 highly variable genes via scanpy. Imaging data were scaled from 16-bit to unit range, contrast-adjusted per channel using percentile clipping, and resampled with bilinear interpolation. Canonical SMILES for PubChem drugs were obtained with up to 20 variants enumerated to train molecular autoencoders; evaluation outputs were standardized into canonical SMILES with RDKit Landrum (2016).

For single-modality experiments, we further incorporated preprocessed LINCS gene expression data Subramanian et al. (2017) covering *>*14,000 perturbations with matched PubChem records. To mitigate LINCS batch effects, we applied Harmony Korsunsky et al. (2019) for dimension-matched normalization. We also evaluated on CPG-jump Cell Painting data Chandrasekaran et al. (2024), comprising ∼ 300 compounds across two cell lines (U2OS and A549), for imaging-only experiments. Unlike GDP, the model was not trained on any CPG-jump or LINCS data; both datasets draw on substantially larger compound libraries than GDP, and all compounds evaluated in these two datasets are unseen by the model, so CPG-jump and LINCS results reflect genuine zero-shot generalization to structurally novel drugs.

### 5.6 Evaluation Metrics

We evaluate the quality of generated molecules using a diverse set of metrics. *Validity* measures the percentage of chemically valid SMILES strings. *Fréchet ChemNet Distance* (FCD) quantifies the distributional distance between generated molecules and the training reference set using activations from a pre-trained chemical neural network (evaluated under training-set conditions). *Quantitative Estimate of Drug-likeness* (QED) provides a unified drug-likeness score, while *Lipinski compliance* tracks adherence to the Rule of Five (MW *<* 500, LogP *<* 5, H-bond donors *<* 5, acceptors *<* 10). We assess therapeutic relevance using *Tanimoto Similarity*, calculated on Morgan fingerprints (radius 2, 2048 bits) between generated molecules and ground-truth reference compounds. To detect mode collapse, we measure *Scaffold Diversity* and compute KL divergences for molecular weight, LogP, and TPSA distributions relative to the training data. Sampling is deterministic as rectified flow integrates a fixed velocity field by Euler steps, so for a fixed initial noise seed the generated set is reproduced exactly across runs. All reported generation metrics therefore use a single fixed seed, and the FCD, QED, and KL values are exact for that seed rather than estimates over stochastic draws.

### 5.7 Experimental Tasks

#### Generation task

Pert2Mol demonstrates the capacity for inverse molecular design by generating molecular hypotheses directly conditioned on biological perturbation profiles. The model employs a rectified flow-based generative process to construct molecular structures along straight-line trajectories, conditioned on encoded control and treatment phenotypic features. This capability directly measures Pert2Mol’s capacity to recover bioactive scaffolds and explore chemically meaningful regions of molecular space while maintaining therapeutically relevant properties.

#### Retrieval task

To validate the precision of the learned mapping on seen data, we perform retrieval accuracy analysis on the augmented training set. Each training sample is encoded to predict a molecular embedding, which is used as a query to search for its exact ground-truth compound within the pre-computed drug latent space of the training set. Retrieval performance is measured using Precision@K and Hit@K (*K*=1, 3, 5, 10) and Mean Reciprocal Rank (MRR). This task verifies that the multi-modal encoder has successfully learned to disentangle compound-specific signatures from the training inputs and map them to the correct region of the chemical manifold.

#### Repurposing task

To evaluate generalization to unseen experimental conditions, we encode test-set biological profiles to produce a predicted molecular embedding, then perform nearest-neighbor retrieval in the pre-computed drug latent space to identify the closest existing therapeutic agents. Unlike retrieval (which operates on training data), repurposing evaluates whether the learned biological-to-chemical mapping generalizes across conditions; because GDP’s train and test partitions share the same 39 compounds (Section 5.5), this task tests generalization to unseen cell line/dose/timepoint combinations of known compounds rather than to structurally novel drugs. Performance is quantified using molecular validity, physicochemical properties (QED, Lipinski), and Tanimoto similarity to the ground-truth drug.

## Supplementary information

Not applicable.

## Declarations

### Funding

### Competing interests

The authors declare no competing interests.

### Ethics approval and consent to participate

Not applicable.

### Consent for publication

Not applicable.

### Data availability

The GDP dataset is available from Ginkgo Bioworks. The CPG-jump dataset is available at Chandrasekaran et al. (2024). The LINCS L1000 dataset is available at Subramanian et al. (2017).

### Code availability

Code and pretrained models will be made available at https://github.com/wangmengbo/Pert2Mol.

### Author contributions

M.W. and S.V. designed the model architecture and conducted experiments. V.M.P. contributed to data processing and evaluation. S.J. and S.K ran the baseline model experiments. M.K. A.G. and N.A.L. supervised the project and provided scientific and biological expertise. All authors read and approved the final manuscript.

## Notes

### Competing Interest Statement

The authors have declared no competing interest.

### Summary of Updates

New results for evaluation. Improved Figures. New authors as scope of project has changed.

